# Neural energy metabolism encodes directed communication across the human cortex

**DOI:** 10.64898/2026.05.28.728560

**Authors:** Roman Belenya, Samira Epp, Antonia Bose, André Hechler, Laura Fraticelli, Mahnaz Ashrafi, Andreas Ranft, Igor Yakushev, Katarzyna Kurcyus, Gabriel Castrillon, Valentin Riedl

**Affiliations:** Department of Neuroradiology, Klinikum rechts der Isar, TUM School of Medicine and Health, Technical University of Munich, Munich, Germany; Department of Neuroradiology, Uniklinikum Erlangen, Friedrich-Alexander-University Erlangen-Nuremberg, Erlangen, Germany; Graduate School of Systemic Neurosciences, Ludwig-Maximilians-Universität München, Munich, Germany; Research Group in Medical Imaging, SURA Ayudas Diagnósticas, Medellin, Colombia; Department of Anesthesiology and Intensive Care Medicine, Klinikum rechts der Isar, TUM School of Medicine and Health, Technical University of Munich, Munich, Germany; Department of Nuclear Medicine, Klinikum rechts der Isar, TUM School of Medicine and Health, Technical University of Munich, Munich, Germany

**Author notes:** Shared contribution (Gabriel Castrillon, Valentin Riedl).

**Keywords:** effective connectivity, functional connectivity, glucose metabolism, fMRI, FDG PET

## Abstract

Complex cognition depends on directed communication between brain regions. Yet, resting functional MRI resolves only undirected associations, and inferring the direction of signalling has proven difficult across the whole brain. Because synaptic transmission is metabolically asymmetric, with the post-synaptic neuron bearing most of the energetic cost, regional energy demand carries a physical signature of the direction of communication. Here, we show that this signature organizes the human cortex into a reproducible directed hierarchy, in which sensory and attentional systems preferentially drive higher-order networks, while the default mode and control networks act as principal receivers. Critically, the inferred directionality is independently predicted by two cellular markers of neuronal energy demand, regional mitochondrial density and laminar cytoarchitecture, linking macroscale information flow to its cellular substrate. The directed architecture recapitulates the known feedforward and feedback organization of visual pathways and, using an average CMR_Glc_ template, extends to functional MRI acquired without PET. By grounding communication direction in energy metabolism, Metabolic Connectivity Mapping bridges correlative and effective connectivity and offers a scalable, biologically interpretable framework for mapping directed communication across the human brain.

## Introduction

Complex cognition arises from dynamic interactions among brain regions that continuously exchange information through directed neural signalling. Resting-state functional MRI (rs-fMRI) reliably maps large-scale functional connectivity (FC) across the human brain (Thomas Yeo et al., 2011; Schaefer et al., 2018), but FC is inherently undirected and reveals little about the direction of information flow. Inferring whether one region drives another, termed effective or directed connectivity, is essential for understanding hierarchical communication, yet it remains difficult at whole-brain scale. Established approaches such as Dynamic Causal Modelling (Friston et al., 2003; Razi et al., 2017; Novelli et al., 2024) and Granger causality (Seth et al., 2015; Friston et al., 2013; Roebroeck et al., 2011) infer directionality from the statistical patterns in the fMRI signal itself, but are constrained by the hemodynamic response and become computationally demanding for whole-cortex networks. A complementary strategy is to derive direction not from signal dynamics, but from a physical constraint on the neural circuits that generate them.

Synaptic communication is energetically asymmetric: the postsynaptic neuron bears most of the metabolic cost of signal transmission (Attwell and Laughlin, 2001; Howarth et al., 2012). Regional energy consumption should therefore act as a marker of cumulative afferent input, making energy metabolism a candidate physical signature of signalling direction. Consistent with this premise, the aggregate functional throughput of a region has recently been shown to predict its glucose metabolism (Castrillon et al., 2023a; Klug et al., 2024). Metabolic Connectivity Mapping (MCM) established the feasibility of this asymmetry-idea by combining fMRI-derived coupling with regional [_18_F]FDG-PET glucose metabolism. In a proof-of-concept (Riedl et al., 2016), we tested the model on visual system connectivity, and the model has since been applied to task conditions (Hahn et al., 2020; Klug et al., 2022). Together, these studies show that directionality is recoverable from energy metabolism, yet only within limited circuits. To become a useful tool, MCM must be established across the entire cortex, validated against independent biology, and made universally applicable.

Here we show that neural energy metabolism encodes a coherent, brain-wide architecture of directed communication. We implement MCM as a whole-cortex model that decomposes each functional connection into directed and balanced components based on metabolic asymmetry between regions, deriving directionality by directly comparing inter-regional energy demand rather than local similarity between FC and metabolism. This means directionality no longer depends on how well a region’s connectivity profile matches its metabolic profile, but on which of two connected regions is metabolically more expensive to supply. Because each region’s metabolism is first normalized by its total connectivity, the comparison reflects energy per connection rather than overall metabolic rate, preventing densely connected regions from being classified as receivers by virtue of their connectivity alone. Applied across the human cortex, the model reveals a reproducible signalling hierarchy in which sensory and attentional systems preferentially drive association and control networks, with the default mode and control networks as principal receivers. Crucially, this inferred directionality converges with two independent cellular markers of neuronal energy demand, regional mitochondrial density (Mosharov et al., 2025) and laminar cytoarchitecture reflecting cell density in the input layers (Amunts and Zilles, 2015; Wagstyl et al., 2020), linking macroscale information flow to the metabolic and structural substrate of synaptic input. The model further recapitulates feedforward and feedback organization within the visual streams and, using a population-average CMR_Glc_ template, generalizes to large fMRI datasets acquired without concurrent PET.

Together, these results posit MCM as a physiologically grounded, scalable model of directed cortical communication. By grounding signalling direction in energy metabolism and anchoring it to cellular measures of energy demand, our framework bridges correlation-based FC and biologically constrained effective connectivity, providing a biologically interpretable route to map directed communication across the human brain.

## Results

### The biophysical model of MCM

We implemented MCM to infer directed connectivity across the entire human cortex by integrating functional coupling with regional energy consumption from simultaneous PET/MRI measurements (Figure 1). Our model combines BOLD fMRI-derived FC, restricted to anatomically connected region pairs via structural connectivity (SC), with regional measures of absolute glucose metabolism. For each pair of connected regions, we computed an energy ratio to determine the dominant direction of activity flow, generating two complementary matrices: an undirected connectivity matrix representing metabolically balanced interactions and a directed connectivity matrix capturing asymmetric, energy-dependent signalling. Crucially, the decomposition conserves the total interaction strength, such that directed (D_A→B_) and undirected (U_AB_) interactions sum to the original full connectivity (FC_AB_; see Figure 1 box: Metabolic Connectivity Mapping model).

**Figure 1.**
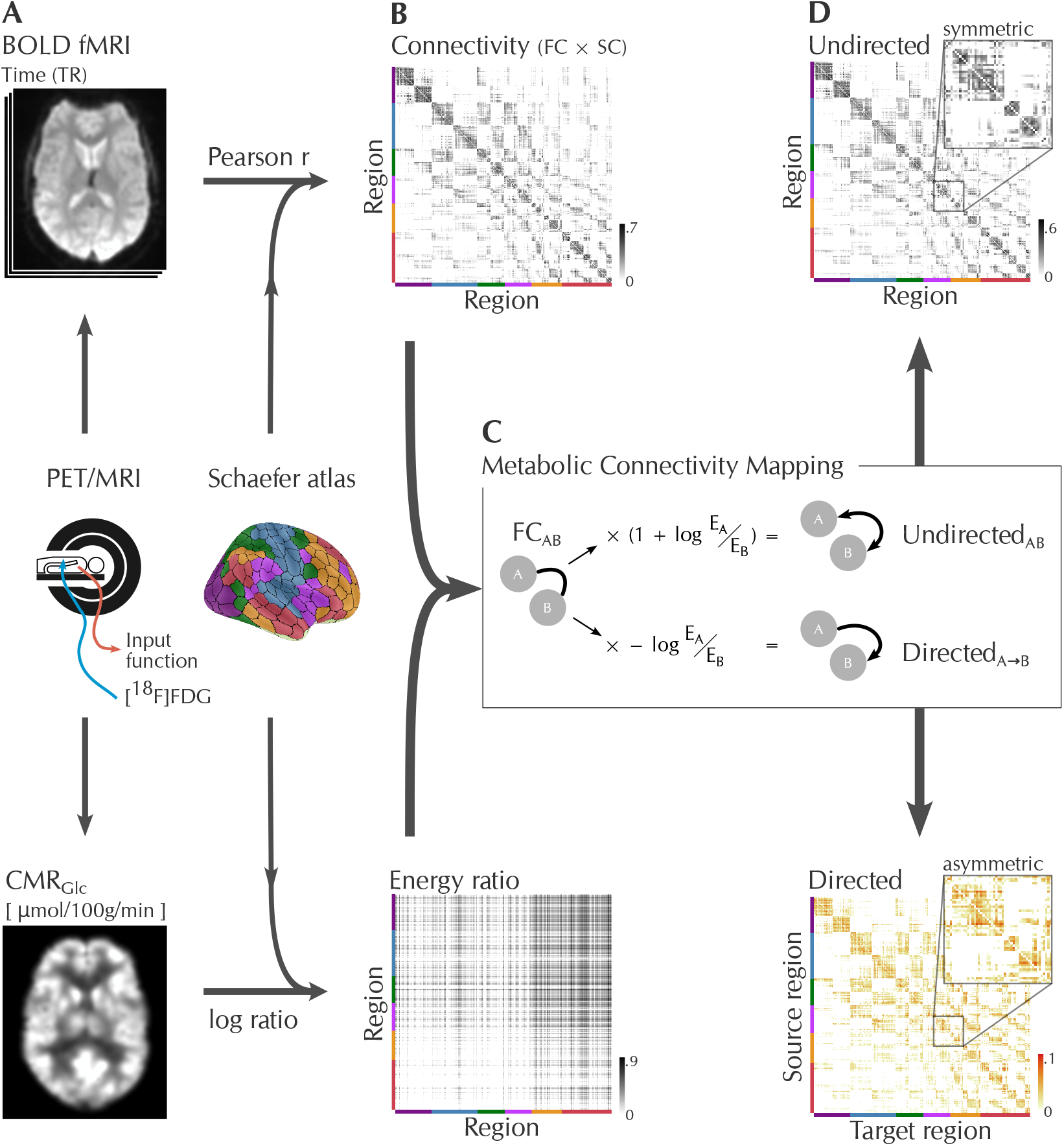
Schematic and implementation of Metabolic Connectivity Mapping (MCM). This figure illustrates the workflow for integrating metabolic and functional imaging to infer directed connectivity. **A**, Simultaneous acquisition of BOLD fMRI (top left) and quantitative [_18_F]FDG PET (bottom left) provides measures of resting-state fMRI signal time series and regional glucose metabolism (CMR_Glc_), respectively. Cortical parcellation divides the brain into 400 regions and associated resting-state networks (Schaefer atlas, coloured surface). **B**, The Connectivity matrix derives from integrating FC and SC, while the Energy ratio matrix is the logarithm of all ratios of the regional energy consumption values. **C**, The box (“Metabolic Connectivity Mapping”) explains the mathematical decomposition of connectivity into directed and undirected components; see Methods for model equations. In brief: When two regions are connected, the region that consumes more energy (E) receives that connection, and the strength of that directed connection is proportional to the region’s energy difference. **D**, Example output: the undirected matrix (top right) captures metabolically balanced connections, while the directed matrix (bottom right) reflects asymmetric, energy-driven information flow. The sum of both matches the original FC.

### Energy asymmetry organizes a cortical signalling hierarchy

Applied across the cortex, MCM assigned a reproducible direction to approximately 14% of all functional connections, a biologically constrained subset that organizes the remaining connectivity into a functional hierarchy (Figure 2A). Figure 2B illustrates regions with the highest proportions of outgoing and incoming connections. Aggregating region-wise directionality into a network-level matrix revealed pronounced asymmetries in network communication (Figure 2C-D). The dorsal attention, ventral attention, and visual networks exhibited the strongest overall outgoing signalling, sending information to other systems more than they received. In contrast, the default mode and control networks acted primarily as receivers, accumulating input while emitting fewer outgoing signals. The somatomotor network displayed a balanced in-out profile.

**Figure 2.**
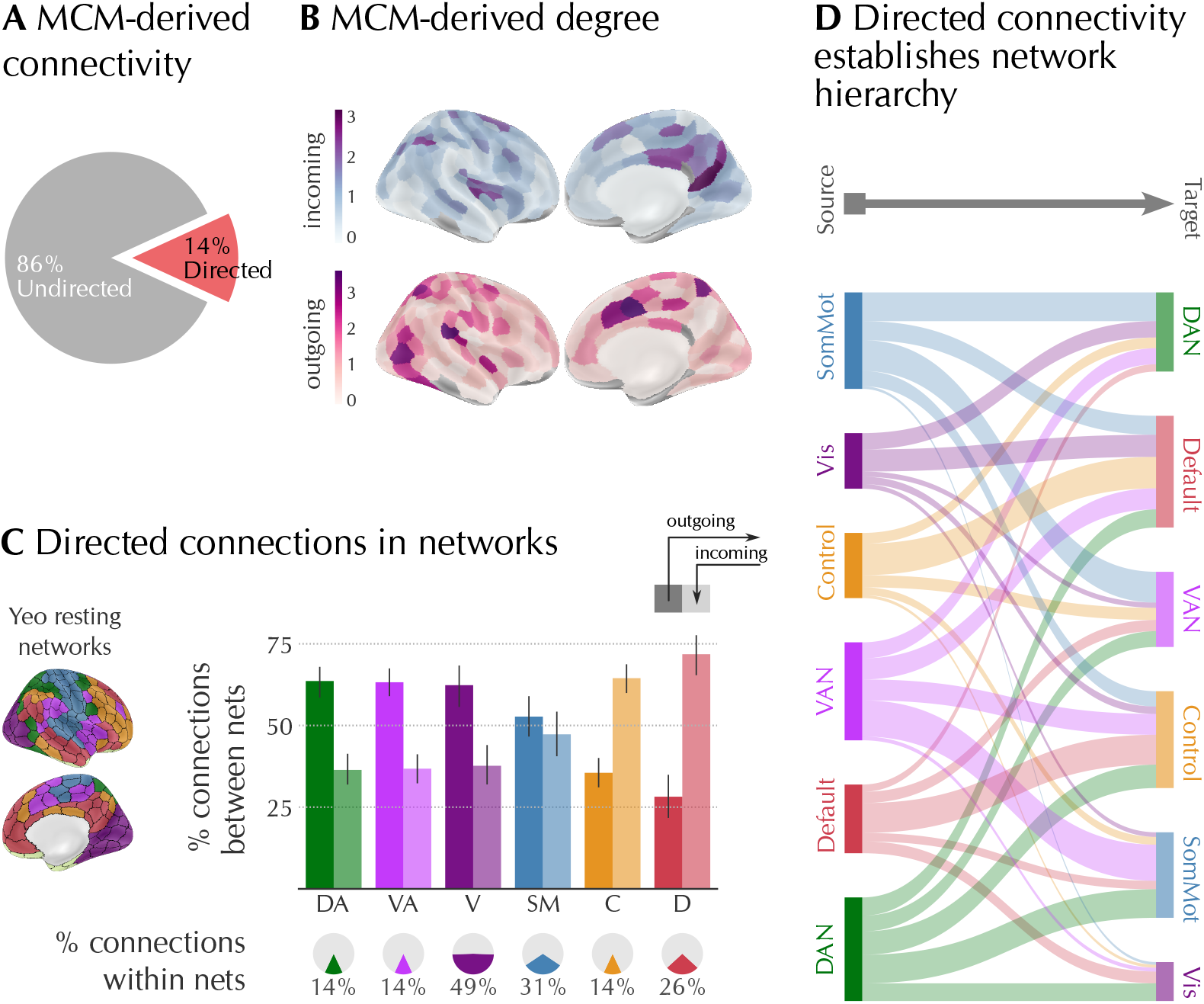
Signalling asymmetries among large-scale brain networks. **A**, Pie chart summarizing the proportion of directed (red) and undirected (grey) connectivity within the cortex as derived from MCM; *≈*14% of the total MCM-derived connectivity is directed. **B**, Surface map displaying regional in-degree (top, blue: receiving) and out-degree (bottom, red: sending) computed from directed connectivity, highlighting hierarchical zones of information flow. **C**, Bar plots summarizing the proportion of outgoing (darker colour) versus incoming (lighter colour) connections for each Yeo network. Bottom: Pie charts represent the fraction of within-network directed connections (i.e. the diagonal of connectivity matrix) for each network, illustrating strong internal integration in the visual and somatomotor networks. **D**, Sankey diagram visualizing directed connectivity between networks; line width represents normalized connection strength. This demonstrates hierarchical asymmetry with the default mode and control networks as principal information receivers and the visual/attention networks as major senders.

Within-network analysis (Figure 2C, pie charts) demonstrated especially strong internally directed connectivity in the visual (49%) and somatomotor (31%) networks, indicating high integration within sensory processing systems. The Sankey diagram (Figure 2D) of the between-network signalling hierarchy revealed the default mode network as the largest information sink, receiving convergent input from attentional and sensory systems. To ensure that results were not biased by network size, we normalized the summed MCM connectivity matrix by the total in- and out-degree of each network (see Methods), allowing for a fair comparison across networks. In summary, cortical directionality followed a hierarchical gradient consistent with known anatomical and functional organization (Margulies et al., 2016; Wang et al., 2019). Sensory and attentional networks displayed predominantly outgoing signalling toward higher-order networks, while the default mode and control networks accumulated convergent afferent input (Figure 2B-D).

### MCM recovers hierarchical processing in the visual streams

To test whether MCM can identify known hierarchical signalling patterns, we analyzed directed connectivity within visual information processing streams. In the early visual cortices (V1, V2 and V3; Figure 3B, “Early”), the strongest edges between complexes were top-down (Figure 3D). At the same time, each area’s total outgoing connectivity to the rest of the visual areas increased along the hierarchy: although the three areas’ outgoing profiles were highly similar (V1 vs V2: r = 0.97; p < 0.001; V1 vs V3: r = 0.91; p < 0.001; V2 vs V3: r = 0.97; p < 0.001; N = 12 complexes), overall outgoing strength was greater for V3 than V2 (t = 5.23, p < 0.001, N = 12 complexes) and V1 (t = 4.87, p < 0.001, N = 12 complexes), and greater for V2 than V1 (t = 3.43, p = 0.006, N = 12 complexes; Figure 3C, Early Stream, rows 1-3). Thus, the dominant “within-triad” edges are feedback, even as system-wide output rises in the bottom-up direction. In the dorsal stream, our model replicated the hypothesized bottom-up hierarchical processing with predominant connections following the V3A → V6/V7 → IP, V3A → MT+→ IP, MT+ → ST connections (Figure 3D, Dorsal stream). However, we found a predominant top-down connection between V3A and V3, linking the dorsal stream to the early visual cortices. Similarly, in the ventral stream, our model replicated the hierarchical processing pattern except for top-down connectivity between FFC and V8 (Figure 3D, Ventral stream). Finally, regions in the IP complex had predominant outgoing connectivity to the ST complex, contrary to the hypothesized direction. Overall, MCM recovered the expected hierarchical processing across the visual streams, with a few exceptions, most notably the feedback dominance in the early visual stream, consistent with the strong modulatory feedback that extrastriate areas exert on early visual cortices.

**Figure 3.**
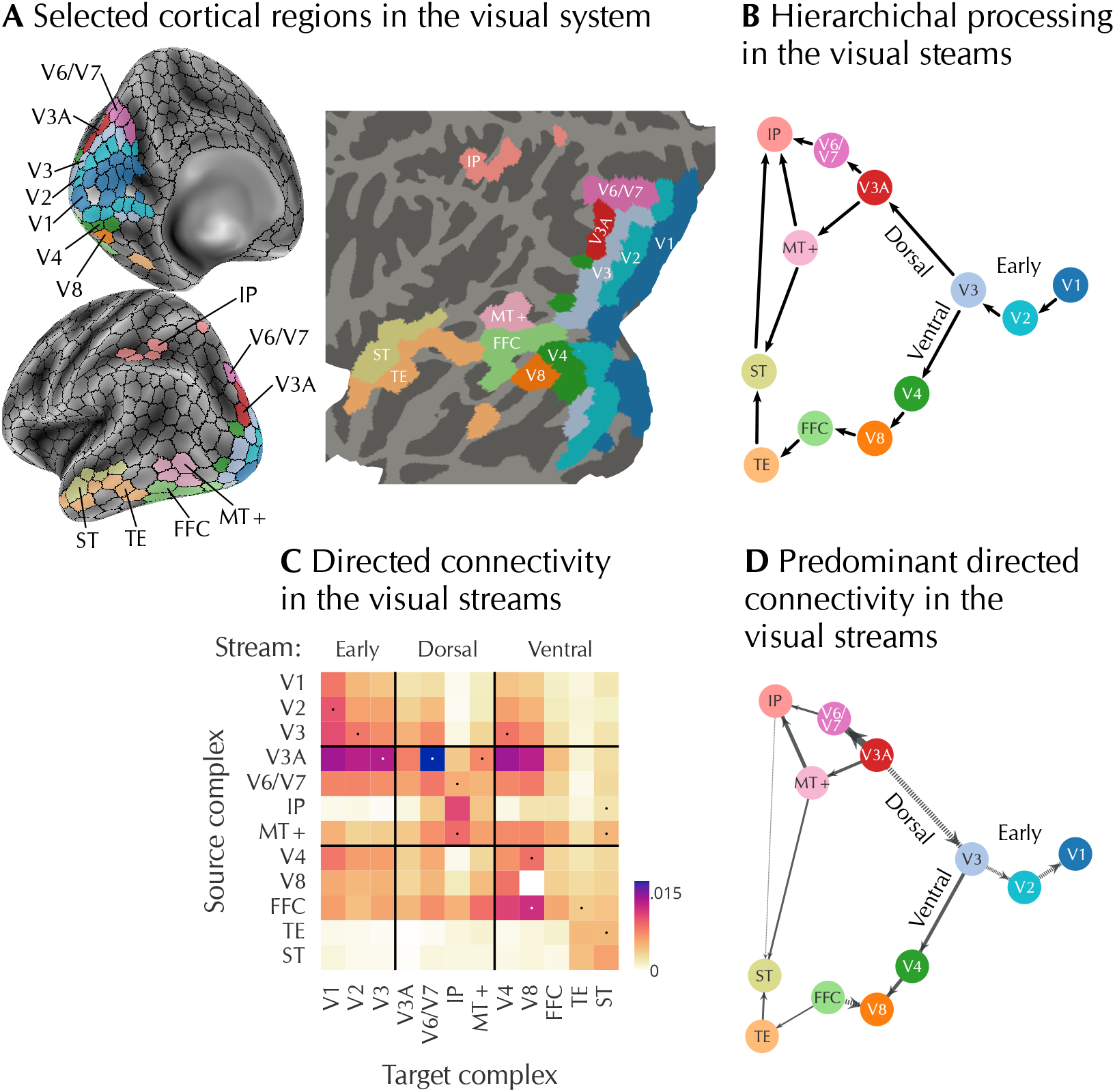
Hierarchical structure and directionality the visual network. **A**, Parcellation of Schaefer regions in the early, dorsal and ventral visual streams on the inflated cortical surface (left) and flattened cortex representation (right). IP, intraparietal complex; FFC, fusiform face complex; MT+, middle temporal complex; TE, inferior temporal complex; ST, medial superior temporal visual complex. **B**, Schematic representation of hierarchical information processing in the visual cortex (adapted and simplified from Ungerleider (1995)). **C**, MCM-derived directed connectivity matrix averaged across N = 20 participants summarizing connectivity strength between regions defined in subfigure A. Connections visualized in subfigure D are marked with a dot. **D**, Network graph showing the predominant connectivity direction for a subset of connections from subfigure D outlined in subfigure B. The arrow width represents the average connectivity strength. Connections with predominant top-down connectivity are plotted with dashed lines.

### Incoming connectivity tracks mitochondrial density

Since mitochondrial density reflects regional energy demand, we tested whether it tracks MCM-derived directionality. The MCM in-degree correlated with mitochondrial density (r = 0.40; p_smash_ < 0.001; N = 362 regions; Figure 4B, bottom left), whereas outgoing degree did not (r = 0.01; p_smash_ = 0.495; N = 362 regions; Figure 4B, bottom right). This dissociation is predicted if incoming connections carry the postsynaptic energy cost. As a validity check on the marker itself, regional CMR_Glc_ and mitochondrial density were also strongly correlated (r = 0.56; p_smash_ < 0.001; N = 362 regions; Figure 4B, top left), confirming that glucose consumption and mitochondrial abundance co-vary across cortex. In a joint regression, incoming and outgoing degrees together explained 17% of the variance in mitochondrial density (R^2^ = 0.172), with in-degree accounting for 95% and out-degree for only 5% of the shared variance. Thus, mitochondrial density aligns with both glucose consumption and incoming connectivity, linking MCM-derived direction to a cellular marker of energy demand.

**Figure 4.**
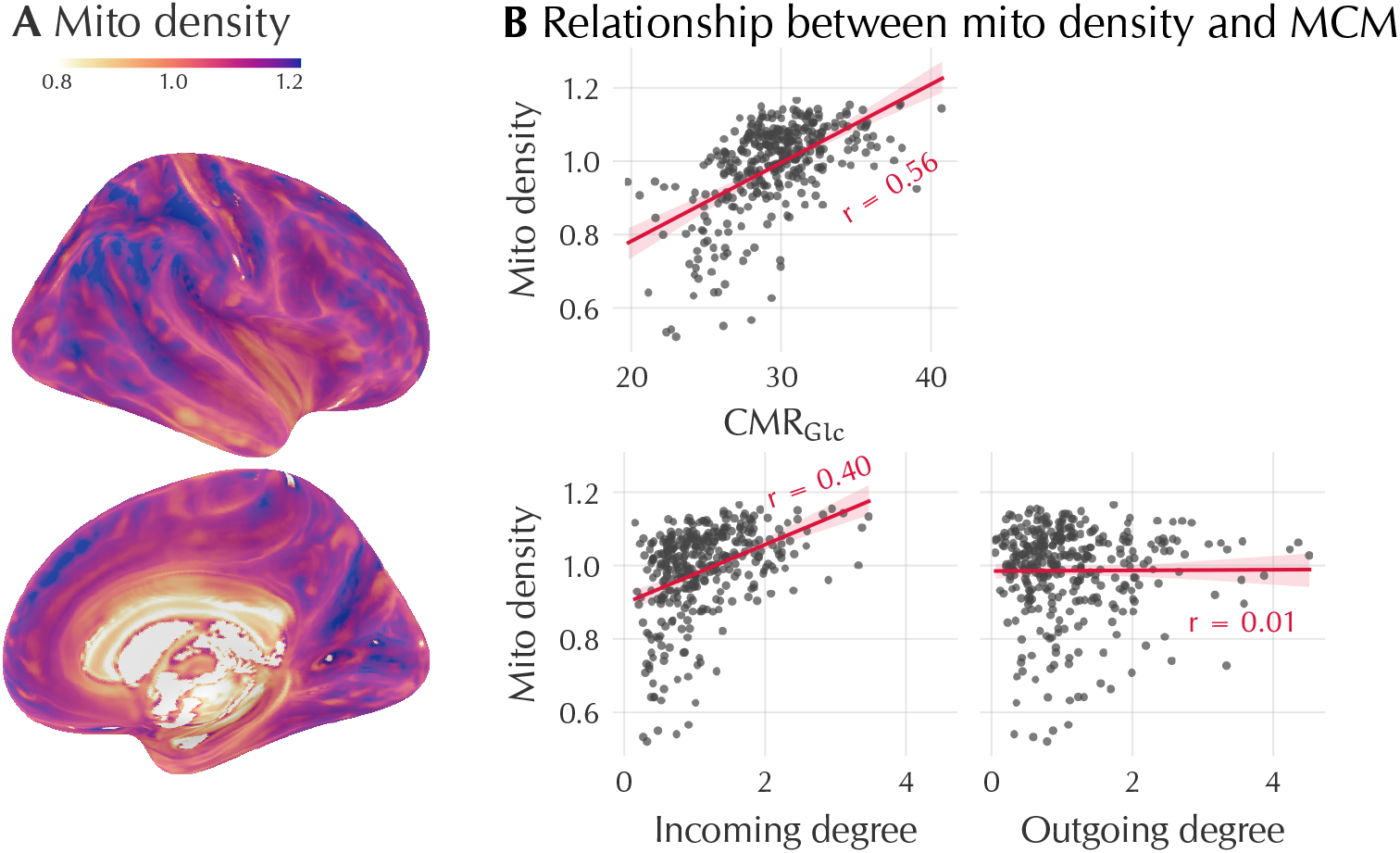
Correlation of metabolic directionality with regional mitochondrial density and cortical cytoarchitecture. **A**, inflated map of the right hemisphere depicting predicted regional mitochondrial density (Mosharov et al., 2025), an independent marker of glucose oxidation capacity. **B**, Top left: scatter plot comparing CMR_Glc_ to mitochondrial density, revealing significant linear association (r = 0.56; p_smash_ < 0.001) across 362 regions. Bottom: MCM-derived degree versus mitochondrial density, where only incoming degree shows robust association after controlling for spatial autocorrelation (r = 0.40, p_smash_ < 0.001). Red lines denote linear regression fits; dots, Schaefer regions; shaded area, 95% CI. r denotes Pearson correlation.

### Input regions carry dense superficial layers

We next examined how directionality relates to cortical microstructure using cell density data from the Big-Brain atlas (Amunts et al., 2013). Cortical cytoarchitecture refers to the distribution pattern of the cell bodies across cortical layers. Cells located in superficial layers typically receive projections from other cortical areas, while pyramidal neurons in deeper layers send outputs to distant cortical targets (Amunts and Zilles, 2015). This anatomical organization suggests that the location of cell bodies across the cortical depth can serve as a proxy for incoming or outgoing connectivity. To parametrize the cell density distribution, we computed the average regional cortical depth and skewness from a dataset that sampled the BigBrain at 50 cortical depths (Figure 5A, left; Wagstyl et al. (2020)). In-degree correlated significantly with both the skewness (r = 0.28; p_smash_ = 0.006; N = 362 regions) and the depth (r = −0.30; p_smash_ = 0.002; N = 362 regions) of staining intensity profiles (Figure 5B, left column).

**Figure 5.**
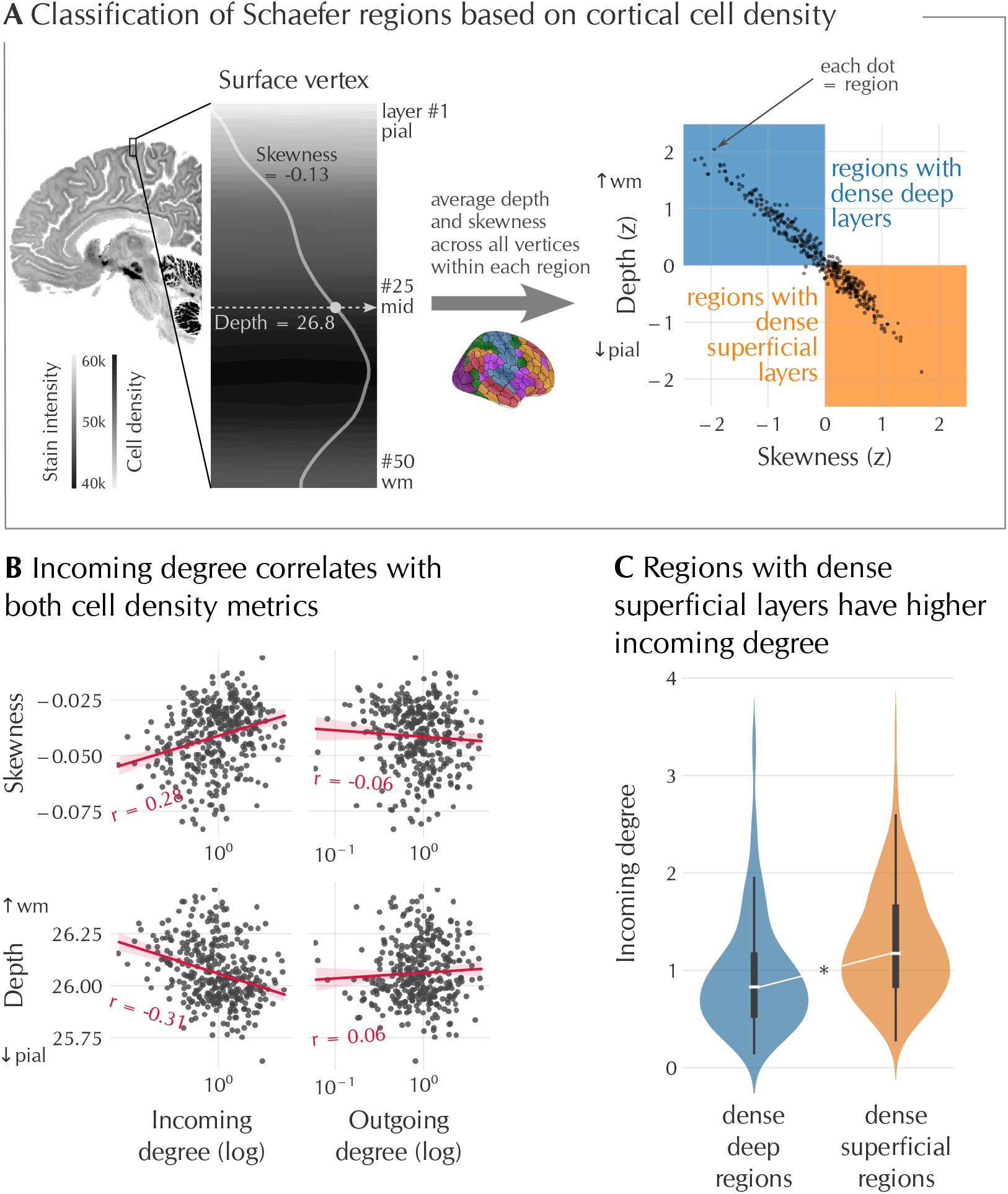
Input regions have higher cell density in the superficial layers **A**, Left: illustration of the laminar staining intensity profile sampled across 50 equivolumetric cortical depths from pial (superficial, top) to white matter surfaces (deep, bottom) for a single surface vertex of the BigBrain template (depicted as a volumetric sagittal slice for simplicity) (Wagstyl et al., 2020). A darker colour represents lower staining intensity and thus higher cell density. Depth is the layer number at which cell density is highest across cortical depth, represented by the weighted average of the distribution (grey line). Skewness represents the location of the distribution tail. Right: binarization of regions into “dense superficial” (orange) and “dense deep” (blue) profiles based on weighted mean cell depth and skewness (see Methods). **B**, Scatter plots: MCM in-degree versus average staining skewness and depth for all regions (left two panels). **C**, Violin plot comparing in-degree for “superficial” vs “deep” regions; regions with denser superficial layers (input recipients) display significantly higher incoming connectivity. All statistics are corrected for spatial autocorrelation.

Because lower depth and more positive skewness both index cell density shifted toward superficial layers (Figure 5A), these associations place high in-degree regions among those with dense superficial profiles, consistent with their role as recipients of cortico-cortical input. Out-degree, by contrast, showed no significant correlation with either skewness (r = −0.06) or depth (r = 0.06), the same in/out dissociation seen for mitochondrial density. To fully characterize the cell location, we combined the depth and skewness parameters as suggested by Palomero-Gallagher and Zilles (2018), thus classifying the 400 Schaefer regions into dense deep and dense superficial profiles (Figure 5A, right). Regions with dense superficial layers showed significantly higher in-degree than dense-deep regions (t = −5.13; p = 0.0002; N = 362 regions), consistent with their role in receiving cortical projections (Figure 5C). In summary, MCM in-degree reflects a defining cytoarchitectonic feature of input cortex: regions with cell density concentrated in the superficial layers preferentially receive cortical projections.

### Template-based MCM generalizes to MRI-only data

To test whether MCM applies to datasets without PET, we computed directionality for the fMRI data from the Human Connectome Project (HCP Young Adult 100 Unrelated subjects, Glasser et al., 2013) by replacing individual CMR_Glc_ maps with our group-average CMR_Glc_-template (Figure 6A).

**Figure 6.**
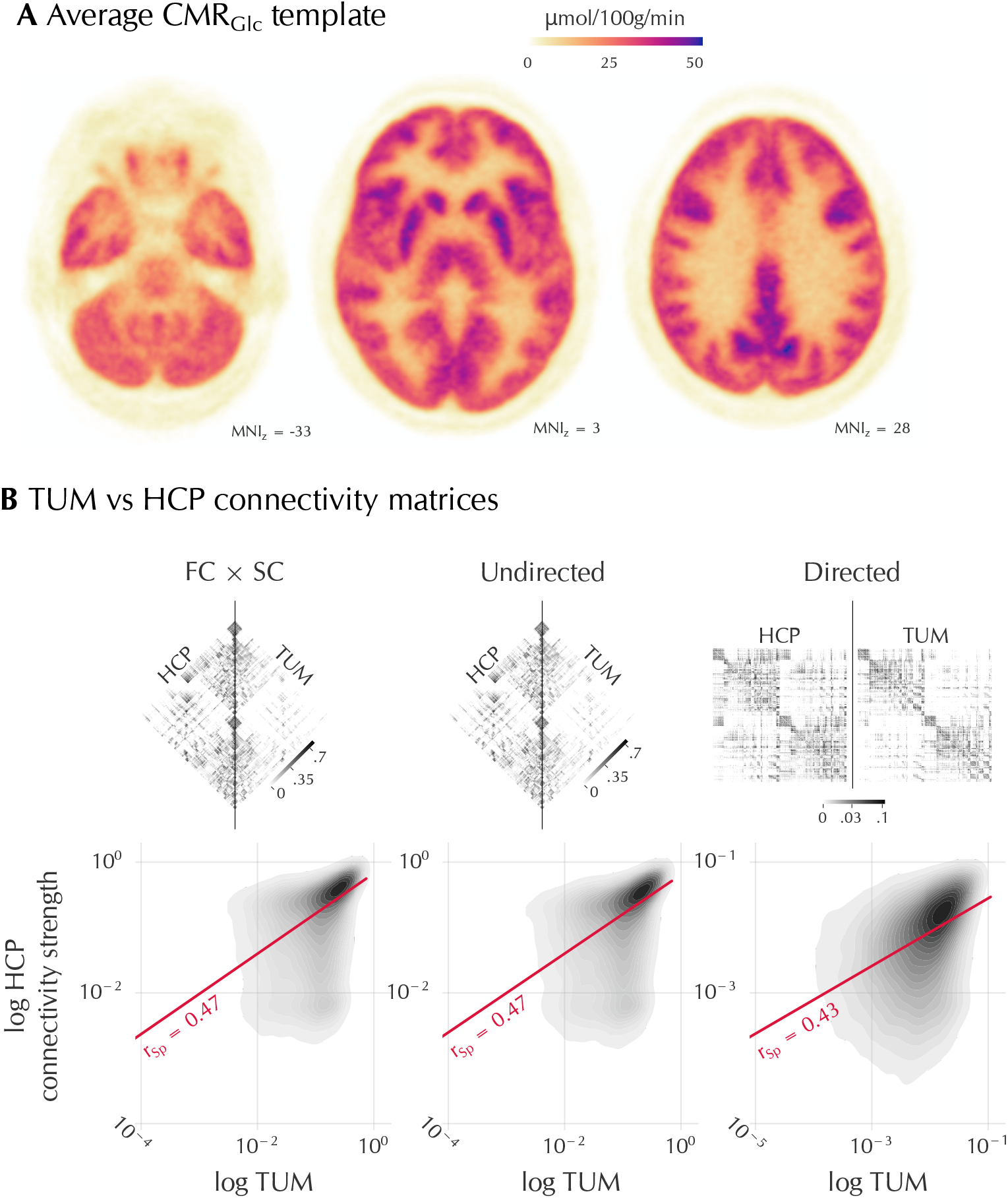
Applicability of the CMR_Glc_ template model to large-scale fMRI datasets. **A**, Axial brain slices visualizing our group-averaged CMR_Glc_ template in standard MNI space, used for template-based MCM calculations. **B**, Connectivity matrices (top row) visualize the group-averaged HCP (fMRI-only plus template) and TUM (derived from PET/fMRI data) connectivity patterns side-by-side. Symmetrical matrices (FC ×SC and Undirected) are split and merged across the diagonal (i.e. the HCP-derived matrix is on the left and the TUM-derived matrix is on the right-hand side). Because directed matrices are asymmetrical, the full HCP and TUM matrices are shown side-by-side. Scatter plots quantify the similarity between the connectivity matrices; FC × SC (left, r = 0.47), undirected (middle, r = 0.47) and directed (right, r = 0.43) components, both Spearman rank correlation (r_Sp_; p < 0.001 using 10,000 null models). Data are log-transformed and visualized using 2D kernel density estimates to account for the power distribution of connectivity values.

Despite differences in preprocessing, HCP-derived connectivity matrix matched the TUM cohort (r = 0.47; p_null_ < 0.001; N = 19252 non-zero connectivity matrix entries, Figure 6B left). Similarly, despite the absence of individual PET, the HCP cohort matched the TUM cohort for both the undirected (r = 0.47; p_null_ < 0.001; N = 19252 non-zero connections) and directed (r = 0.43; p_null_ < 0.001; N = 31595 non-zero connections) components (Figure 6B, centre and right). That the correlations are moderate rather than near-perfect is expected given the fully independent cohort, scanner, preprocessing, and the substitution of individual metabolism with the group template; the preserved rank structure indicates that MCM’s directional organization is carried by the template, not by subject-specific PET. Together, these results show that MCM can be applied retrospectively to large fMRI datasets lacking PET by substituting individual maps with a normative metabolic reference. Because MCM reads directionality from regional metabolism, conditions with focal metabolic alterations, such as Alzheimer’s disease, are a suitable application: the same template substitution allows directional hierarchies to be estimated in existing fMRI cohorts. We release the average CMR_Glc_ template publicly: https://github.com/NeuroenergeticsLab/metabolic-connectivity-mapping/tree/main/cmrglc-templates.

## Discussion

MCM infers the direction of cortical communication from regional energy metabolism. Applied across the whole cortex, it recovers a reproducible directed hierarchy in which sensory and attentional systems act predominantly as drivers, while the default mode and control networks are receivers. This directionality corresponds with regional mitochondrial density and laminar cytoarchitecture, linking the imaging-derived hierarchy to cellular markers of energy demand. Compared with the proof-of-concept (Riedl et al., 2016), which inferred direction from the spatial similarity between a region’s functional connectivity and metabolism, the present model (i) derives all directed connections simultaneously rather than one region at a time, (ii) scales from circumscribed circuits to the whole cortex, (iii) normalizes each region’s metabolism by its total connectivity, so that direction reflects energy per connection rather than overall metabolic rate, and (iv) replaces individual PET with a shareable metabolic template, while retaining the same neuroenergetic basis.

Conceptually, MCM assumes that the postsynaptic energy asymmetry established for single synapses aggregates at the regional scale, so that a region’s overall glucose metabolism reflects the postsynaptic cost of the input it receives, and the higher-consuming region in a connected pair is the receiver. This follows from Attwell’s energy budget for grey matter, in which restoring postsynaptic ion gradients is the dominant term of the signalling cost (Attwell and Laughlin, 2001; Howarth et al., 2012). Because [_18_F]FDG-PET measures total glucose demand rather than a specifically postsynaptic component, the directional signal could instead reflect a nonspecific coupling between metabolism and connectivity. Three of our findings argue against this. First, the association is directionally specific: incoming degree correlates with both mitochondrial density (Figure 4) and laminar input architecture (Figure 6), whereas outgoing degree does not. Second, the premise that regional metabolism reflects synaptic signalling is supported by independent work, both by direct measurements (Lundgaard et al., 2015; Viswanathan and Freeman, 2007), by modelling of functional input (Klug et al., 2024), and by our prior work on integrating neurotransmitter (Castrillon et al., 2023a) and transcriptomic (Ashrafi et al., 2026) data with energy metabolism. Third, glycolytic enzymes are present in the postsynaptic density (Wu et al., 1997) and are dynamically regulated with synaptic plasticity (Marder and Goaillard, 2006). Consistent with this, the regional distribution of aerobic glycolysis on the macroscale resembles that of MCM-derived incoming connectivity (Vaishnavi et al., 2010). Together, these observations indicate that MCM reads directionality from the total energetic cost of receiving input, consistent with the asymmetry it assumes. Beyond multimodal PET/MRI, the model can be applied to fMRI alone through a standardized metabolic template, extending its reach to datasets without concurrent PET.

Only a subset of functional connections (*≈*14%) exhibit clear directional dominance, yet these asymmetric pathways are sufficient to organize the cortex into a structured hierarchy. At rest, information primarily flows from sensory and attentional systems to association and control regions, with the default mode and control networks serving as major information receivers. The visual and somatosensory networks are the most self-directed, indicating a significant internal information processing before the signalling is relayed to the higher-order networks. This organization parallels theoretical descriptions of cortical hierarchy derived from laminar, functional, and gene-expression gradients (Margulies et al., 2016; Wang et al., 2019). The predominance of balanced, bidirectional connectivity across most regions may reflect recurrent processing loops that sustain intrinsic activity and feedback modulation (Semedo et al., 2022; Nashef et al., 2022). However, it may also reflect the limited capacity of macroscale brain imaging to resolve fine-grained circuits. Either way, directional asymmetry appears to be a selective rather than global property of the connectome, concentrating in pathways that mediate cross-hierarchical integration.

Following up on the high degree of internal processing in the Visual network, our model replicated large parts of the well-known processing hierarchies in the dorsal and ventral visual streams (Ungerleider, 1995; Milner and Goodale, 2008). Although most connections between the primary and extrastriate visual cortices are reciprocal (Felleman and Van Essen, 1991), visual information is processed sequentially, with increasing feature complexity in higher-order visual areas. Our result, however, emphasized the dominance of feedback over feedforward connections in the early visual stream, following growing evidence of the top-down influence on V1 (Gilbert and Li, 2013) (REF). Feedback connections from extrastriate cortices to V1 cortex influence feedforward activity, thereby modulating attentional mechanisms, and have been identified in tract-tracing studies in both rat (Johnson and Burkhalter, 1997) and macaque brains (Siu et al., 2021; Markov et al., 2014). Alternatively, the early visual cortices are known to exchange strong connections with subcortical structures such as the lateral geniculate nucleus and the pulvinar (Casanova and Chalupa, 2023), which our model misses at the current imaging resolution. Overall, our findings align with established microcircuit hierarchies, supporting the anatomical interpretability of MCM while highlighting where macroscale metabolism cannot capture subcortical contributions.

Biological validation of the model supports its neuroenergetic assumptions as two independent cellular markers converge on the same directional signal. Incoming connectivity was associated with mitochondrial density, consistent with the elevated metabolic demand of postsynaptic dendrites (Tröger et al., 2025), and with dense superficial layers, the cytoarchitectonic signature of regions that receive corticocortical input (Palomero-Gallagher and Zilles, 2018; Genescu and Garel, 2021). In both cases, the effect was specific to incoming connectivity and survived correction for spatial autocorrelation, so the correspondence is unlikely to arise from shared large-scale gradients alone. These cross-scale links tie macroscale directionality to cellular organization, suggesting that energy demand constrains the direction of cortical information flow.

To facilitate broader adoption, we developed a population-average CMR_Glc_ template and confirmed strong replication on the HCP Young Adult dataset (Glasser et al., 2013). This result highlights MCM’s scalability beyond specialized PET/MRI setups, offering a computationally tractable method for estimating directionality in large cohorts. Since altered brain metabolism is a hallmark of many neurological and psychiatric conditions (Castrillon et al., 2023a; Yang et al., 2021; Vlassenko et al., 2010), template-based MCM could enable retrospective analyses of disruptions to the energetic hierarchy in disease.

Several limitations warrant consideration. The current spatial resolution limits reliable estimation of directionality in small subcortical nuclei, such as the lateral geniculate nucleus (Casanova and Chalupa, 2023), and in other thalamic regions. High-resolution fMRI and next-generation PET scanners may extend MCM’s applicability to these deep structures. Furthermore, while MCM reflects metabolic demand, directionality may also depend on context-dependent neuromodulatory states or oscillatory phase relationships that are not captured here (Bazinet et al., 2023). Incorporating these physiological dimensions could refine estimates of dynamic signalling direction in future research.

In summary, MCM grounds the inference of directed connectivity in energy metabolism, recovering a cortical hierarchy that aligns with cellular and laminar architecture and generalizes to fMRI datasets without concurrent PET. By linking connectivity to the brain’s energetic organization, it offers a biologically interpretable and scalable approach to mapping directed communication in the human brain.

## Methods

### Dataset

For a full description of the data acquisition, refer to (Castrillon et al., 2023a). Briefly, our dataset consisted of twenty BOLD fMRI and [_18_F]FDG PET recordings from 10 female and 10 male participants (mean age = 35 ± 10 years). An integrated 3T PET/MRI Siemens Biograph scanner (Siemens, Erlangen, Germany) simultaneously acquired the data. Participants received a bolus intravenous injection of [_18_F]FDG (average dose = 184 ± 8 MBq), and the Twilight blood counter measured the arterial input function (AIF) by drawing the arterial blood samples every second throughout the acquisition. Participants were instructed to keep their eyes open throughout the scanning session. Additionally, we had T1-weighted MPRAGE (voxel size = 1 mm) and DWI (voxel size = 2 mm, 30 directions at b = 800 s/mm_2_ and one b = 0 s/mm_2_ volume in AP and PA phase-encoding directions) scans.

### Data preprocessing

#### Structural images

The `antsCorticalThickness` pipeline from the ANTs package preprocessed the T1w images. The processing steps included bias field correction, image denoising, brain extraction and tissue-type segmentation (Avants et al., 2011). In addition, the FreeSurfer `recon-all` pipeline (Dale et al., 1999) generated the pial and white matter surfaces. We aligned the preprocessed images with the MNI152 T1w atlas (MNI 152NLlin6Asym, included in FSL) using the affine (12 DOF) and nonlinear SyN transformations implemented in ANTs.

#### fMRI images

The Configurable Pipeline for the Analysis of Connectomes (CPAC, version 1.4.0, Cameron et al. (2013)) package preprocessed the functional data. Briefly, the images underwent slice-time correction, motion correction, intensity normalization and nuisance regression. Finally, the pipeline removed the low and high frequencies from the time series using a band-pass filter (0.01-0.1 Hz).

We then aligned the preprocessed time-series average to the anatomical images using boundary-based registration with the FSL `flirt` command (Jenkinson and Smith, 2001). We used the resulting transformation matrix to align the cortical parcellation to functional space, thus preserving the functional data in its original state.

We averaged the time series across voxels belonging to the parcellation regions. Then, we computed the functional connectivity (FC) matrices using Pearson correlation of the regions’ average time series. This resulted in a 362 × 362 matrix, and we thresholded the matrices below zero to include only positive correlations in our FC.

#### DWI images

We ran preprocessing and probabilistic white matter tractography using the mrtrix3 suite (Tournier et al., 2019). The preprocessing steps included image denoising, motion and eddy current correction using FSL commands (`topup` and `eddy`), and bias field correction using ANTs. The anatomically constrained tractography (ACT) pipeline estimated response functions using the Tournier algorithm and computed fibre orientation distributions via single-tissue constrained spherical deconvolution. We aligned the preprocessed b0 volume to the anatomical scan using boundary-based registration in FSL’s `flirt`.

The pipeline then estimated streamlines using 5 tissue-type segmentations and the grey-to-white matter interface to constrain the tractography. 20 million tracts were estimated, and a scaling factor was assigned to each streamline with the `sift2` algorithm. To ensure that the streamline weights are proportional to the units of fibre density similarly across all subjects, we further multiplied the scaling factor by the µ estimate from the tractography.

Finally, we constructed the region-wise connectome using the parcellation aligned to the DWI space. The elements in the resulting structural connectivity (SC) matrix were proportional to the white matter bundle cross-sectional area (Smith et al., 2020).

#### [_18_F]FDG PET images

NiftyPET python library reconstructed the raw data from PET recordings offline based on the OSEM algorithm (Markiewicz et al., 2018) (33 frames: 10 frames × 12 s, 8 × 30 s, 8 × 60 s, 2 × 180 s and 5 × 300 s). We corrected the images for attenuation and motion and spatially smoothed the time series with a Gaussian filter of FWHM = 6 mm. We quantified the PET data using the Patlak plot approach (Patlak et al., 1983). The Feng three-exponential function (Feng et al., 1993) estimated the fit to the blood radioactivity data. We then entered the last five frames (5 min each) into a linear regression model, where the integral of plasma radioactivity, adjusted by plasma radioactivity (Patlak x-axis), was the regressor, and tissue radioactivity, adjusted by plasma radioactivity (Patlak y-axis), was the dependent variable. The slope of the regression line (K_i_) is proportional to the [_18_F]FDG intake. Adjusting the slope value with a subject’s glucose concentration in blood, the Lumped constant (Wu, 2003) and grey matter density resulted in the measure of the cerebral metabolic rate of glucose consumption (CMR_Glc_). We further performed partial volume correction on the CMR_Glc_ maps using the Yang algorithm (Yang et al., 2017) and downsampled the maps to an isotropic voxel size of 3 mm.

We aligned the CMR_Glc_ maps with the subjects’ anatomical scans using a rigid-body transform with 6 degrees of freedom (ANTs). The inverse of the resulting transformation matrix aligned the parcellation atlas with the CMR_Glc_ maps in native image space via the anatomical T1w.

#### Cortical parcellation

Schaefer et al. (2018) cortical parcellation with 400 parcels and 7 resting-state (Thomas Yeo et al., 2011) networks defined the brain regions for this analysis. We converted the parcellation into each modality using first, nonlinear and affine transforms between the MNI152 atlas and each subject’s T1w image and second, the affine matrices to align the parcellation with each modality: fMRI, DWI or CMR_Glc_. We performed alignment and nearest-neighbour resampling of the parcellation using the ANTs toolbox in a single step to minimize loss of data precision.

We excluded the entire Limbic network (26 regions) due to the BOLD fMRI signal drop-out in the inferior frontal and temporal lobes. Further, we included a region in the analysis only if it contained at least 5 voxels in both the fMRI and CMR_Glc_ images across all subjects. If a region lacked sufficient voxels in at least one subject, we removed it from all subjects. We identified 38 regions with insufficient data, out of which 22 regions were part of the Limbic network. Thus, we entered 362 regions in the analysis.

### Analysis

#### Average Yeo network connectivity

To assess the connectivity of Yeo networks, we aggregated the region × region (362 × 362) connectivity matrices into the network × network (6 × 6) matrices and calculated several metrics. We first summed all connectivity weights across each network combination. Then, we calculated the indegree and outdegree of this reduced matrix by summing along the rows and columns, respectively. The percentage of in- and outdegree is the in-or out-degree divided by the total degree (incoming + outgoing, excluding the diagonal elements) of the network connectivity matrix. We then computed each network’s self-connections. In the network sums matrix, the diagonal elements represent the total connectivity that originates and terminates within one network. We calculated the percentage of self-connections by dividing the sum of the diagonal elements by the sum of the in- and outdegrees, including the diagonal elements.

Each Yeo network consists of a different number of regions. Thus, to compute a network ×network connectivity matrix adjusted for each network’s size, we divided the network sums by the sum of in- and out-degrees for each network’s combination. For example, for a directed connection between networks A and B, the adjusted connectivity weight was the total connectivity weight connecting A to B, out of the sum of A’s efferent connectivity and B’s efferent connectivity.

#### Analysis of visual streams

To analyze directed connectivity patterns in the visual streams, we first assigned anatomical labels from the HCP MMP parcellation (Glasser et al., 2016) to the Schaefer parcellation. For this analysis, we used the Schaefer parcellation with 500 regions per hemisphere (Schaefer1000, Schaefer et al., 2018) to improve the precision of anatomical definitions. We overlapped the two parcellations in the high-resolution fsaverage space (163K nodes resolution). The mode (i.e. most common value) across all MMP labels within a Schaefer region’s vertices assigned a Glasser anatomical description to each region within the Schaefer parcellation. We further grouped several regions into either the dorsal or ventral streams based on the Glasser et al. (2016) description in the supplementary neuroanatomical results, thereby replicating and simplifying the cortical wiring diagrams outlined in Ungerleider (1995) and Nieuwenhuys et al. (2008). Table S1 shows the assignment of MMP regions to each complex. We then calculated the directed connectivity matrices for each subject, as in our main analysis. We further averaged the connections within each complex in the visual streams and across all subjects to create a complex × complex connectivity matrix (see Figure 3C). To test whether our model follows the known hierarchical processing, we selected a set of connections that follow the bottom-up processing flow (outlined in Figure 3B). We identified the predominant connectivity direction by taking the strongest connection among the two connecting a pair of regions.

#### Statistics and null models

To compare connectivity matrices, we randomly permuted the matrices while preserving each node’s degree distribution using the functions null\_model\_dir\_sign and null\_model\_und\_sign for directed and undirected connectivity, respectively, from the Python implementation of the BCT toolbox (Rubinov et al., 2009). For each connectivity matrix, we calculated 10 000 null models. Spearman correlation estimated the similarity between any pair of connectivity matrices. For symmetric matrices, we compared the upper triangles; for asymmetric matrices, we used all elements except the diagonal.

For the statistical comparison of parcellated brain maps, we used the `brainsmash` Python package (Burt et al., 2020). The program created 1000 permuted surrogate maps that preserve spatial autocorrelation along the brain’s surface. The afni program `SufDist` computed the geodesic distance between all regions within each hemisphere of the Schaefer parcellation. We used the `fsaverage5` surface for the distance calculations and computed distances within each hemisphere separately, as the inter-hemispheric geodesic distance is not defined. For any pair of brain maps a and b, we computed the true Pearson correlation between a and b and compared it with the distribution of correlations between a and 1000 surrogate maps of b. We ran this analysis separately for the left and right hemispheres and reported the largest p-value across the two, assuming that regions in different hemispheres lack spatial autocorrelation.

### MCM Model

Our MCM implementation consists of two inputs. The first is connectivity, which estimates connections between brain regions but lacks directional information. The second is the pairwise energy ratios (normalized to the regions’ degree of connectivity), which inform the model about signalling directionality but lack information on connectivity. Our method derives the directionality of connected brain regions by combining these two components.

#### Connectivity

We used a combination of FC and SC to achieve robust, precise measurement of the connection strength between brain regions. We prioritized anatomical connections while preserving the FC connection weight by multiplying the FC matrix by the thresholded and binarized SC matrix within each subject. The resulting measurements had attributes of both FC and SC: weighted connectivity strength stemmed from the FC measurement, and the presence or absence of anatomical connection from the SC. We called this metric C for combined connectivity, and C_AB_ denotes connectivity strength between regions A and B.

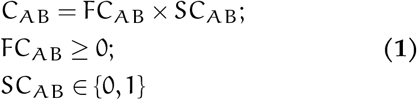

#### Energy ratio

While the CMR_Glc_ data measure regional glucose consumption, our goal was to estimate differences in regional energy use, since our model assumes that the postsynaptic neuron consumes more energy than the presynaptic neuron. We also aimed to estimate energy consumption per connection to allow for all pairwise comparisons. Thus, we first divided each region’s average CMR_Glc_ by the weighted degree of its FC. This metric reflected the amount of energy available per connection within a region. We then divided each region’s edge-wise CMR_Glc_ by that of each other region. Similar to the connectivity matrix, this resulted in a 362 × 362 matrix. We further log-transformed the ratios to scale and center the distribution at zero. After the log-transform, all pairs of regions where the denominator was larger than the numerator were negative and vice versa. We then subset only the negative values because they indicated the afferent direction with respect to the region in the denominator. We inverted the negative values and scaled them to the range [0, 1]. We interpreted the energy ratio as a measure of the directedness of the connection between two regions. We denoted the energy ratio as E and E_AB_ is the ratio of CMR_Glc_ values between regions A and B:

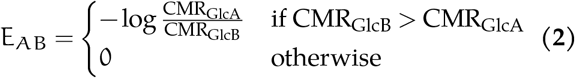

#### Metabolic Connectivity Mapping

Our implementation assumes that the receiving region consumes more energy per connection than the region sensing signalling. Thus, we mapped signalling directionality proportionally to the energy difference for each connected pair of regions. To find directed connectivity, we multiplied the combined connectivity by the energy ratio. This step ensured that directionality was assigned only to the connected regions:

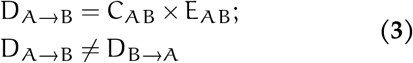

where D_A→B_ is the strength of the directed connection from A to B. Directed connectivity is an asymmetric matrix, where the asymmetry stems from the energy ratio matrix. The opposite of E_AB_ then gives the strength of the undirected connection between A and B, since the values range between 0 and 1:

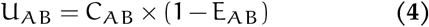

Thus, for two regions A and B, we estimate how directed (D_A→B_), and how undirected (U_AB_) a connection between them is by splitting the functional connectivity into the two components based on their energy differences. For any connection, either D_A→B_ or D_B→A_ was zero (because we removed the ratios where the denominator is less than the numerator, (see Energy ratio section), and the original estimate of connectivity is then the sum of directed and undirected components:

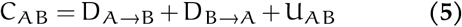

### Average CMR_Glc_ map

To make the MCM analysis applicable to other connectivity studies, we created an average CMR_Glc_ map in the standard MNI152 space (MNI 152Nlin6Asym included in FSL, Evans et al., 2012). The preprocessing steps differed from the subject-specific analysis in several ways. First, from our original 20 datasets, we selected 12 (8 female and 4 male participants, mean age = 32 ± 9 years) with complete AIFs to avoid potential inaccuracies introduced by the average AIF. We aimed to preserve as much data as possible, avoiding unnecessary data resampling and smoothing. We only performed motion correction on the reconstructed [_18_F]FDG PET data and kept the original data resolution (isotropic 1 mm_3_). Patlak plot quantified these data into CMR_Glc_ images.

We then aligned the CMR_Glc_ images with the standard MNI152 space (MNI 152Nlin6Asym) using a rigid-body transform between the PET and T1w spaces and a combination of affine and SyN nonlinear transform between T1w and MNI spaces. The ANTs toolbox performed all image registrations. We computed the average CMR_Glc_ map across all subjects’ maps aligned to the MNI space. In addition, we calculated the CMR_Glc_ map in a newer MNI template (MNI 152Nlin2009cAsym, Fonov et al., 2009) using similar steps, and in the `fsaverage` space (10K and 163K vertex densities) using the freesurfer anatomical preprocessing. The templates, along with the code used in this study, are available on Github: https://github.com/NeuroenergeticsLab/metabolic-connectivity-mapping.

### External datasets

#### HCP

We tested the replicability of our model by applying it to 100 unrelated subjects from the Human Connectome Project Young Adult dataset (Glasser et al., 2013). We downloaded the minimally preprocessed fMRI, DWI and T1w images. For the fMRI images, we resampled the Schaefer parcellation and the grey matter segmentation to a voxel size of 2 mm and computed time-series averages for each region. The rest of the processing pipeline was similar to our own TUM cohort. Since the HCP already distributes the DWI data in anatomical space, we computed only the affine and nonlinear SyN transforms between the MNI and individual T1w images. We applied the transform to the Schaefer parcellation and performed further analysis in the anatomical space. We ran tractography in FSL `probtrackx2` using standard options. This resulted in a region × region SC matrix, which we thresholded and binarized and used to subset connections in the FC matrix. We applied our average CMR_Glc_ map in MNI space to estimate the regional energy ratio, similar to the main pipeline. To compare directed and undirected connectivity with the TUM dataset, we vectorized the matrices and computed Pearson correlations for the non-zero entries.

#### BigBrain

The BigBrain data provide a 3D volumetric atlas of cell-staining intensity from a single post-mortem brain (Amunts et al., 2013). Wagstyl et al. (2020) sampled this atlas in fsaverage space across 50 equivolumetric cortical depths (50 depths × 327K nodes in both hemispheres). In this dataset, the values represent staining intensity, with lower values indicating higher staining intensity (i.e., darker colour). We inverted these data (^1^⁄_x_), so that higher values are proportional to higher cell density. We then calculated the average depth and skewness of the layer-wise intensity distributions. Treating each depth profile as a frequency distribution, we calculated the average depth as the weighted average of cortical depth indices (1-50), with weights inversely proportional to staining intensity. This metric indicated where, across cortical depth, the highest cell density was located. We calculated the skewness as the normalized third moment of the frequency distribution. Skewness measured the location of the distribution’s tail.

We applied the Schaefer parcellation in fsaverage space and computed the average depth and skewness across the vertices belonging to each region. We then classified the brain regions into dense deep and dense superficial profiles, as in (Palomero-Gallagher and Zilles, 2018). The dense deep regions had deeper average depth and more negative skew, while the dense superficial regions had lower average depth and higher average skewness.

#### Mitochondria density

We used the predicted whole-brain mitochondria density map (MitoD) from Mosharov et al. (2025). This map measures mitochondrial density by aggregating cellular markers of the organelle: citrate synthase and mitochondrial DNA. We calculated region-average mitochondrial density using the Schaefer parcellation, as in all other analyses.

## ACKNOWLEDGEMENTS

Valentin Riedl was funded by the European Research Council under the European Union’s Horizon 2020 research and innovation program (ERC Starting Grant, ID 759659).

## DATA AND CODE AVAILBILITY

Raw and preprocessed data are available at OpenNeuro (Castrillon et al., 2023b). The average CMR_Glc_ templates and code we used in this work are available on Github: https://github.com/NeuroenergeticsLab/metabolic-connectivity-mapping.

## AUTHOR CONTRIBUTIONS

RB, conceptualization, data analysis, programming, manuscript writing; SE, data acquisition; AB, data acquisition; AH, programming; LF, data analysis; MA, data analysis; AR, data acquisition; IY, data acquisition; KK, data acquisition; GC, manuscript writing, supervision; VR, conceptualization, manuscript writing, supervision, funding.

## Supplementary Information

**Table S1.**
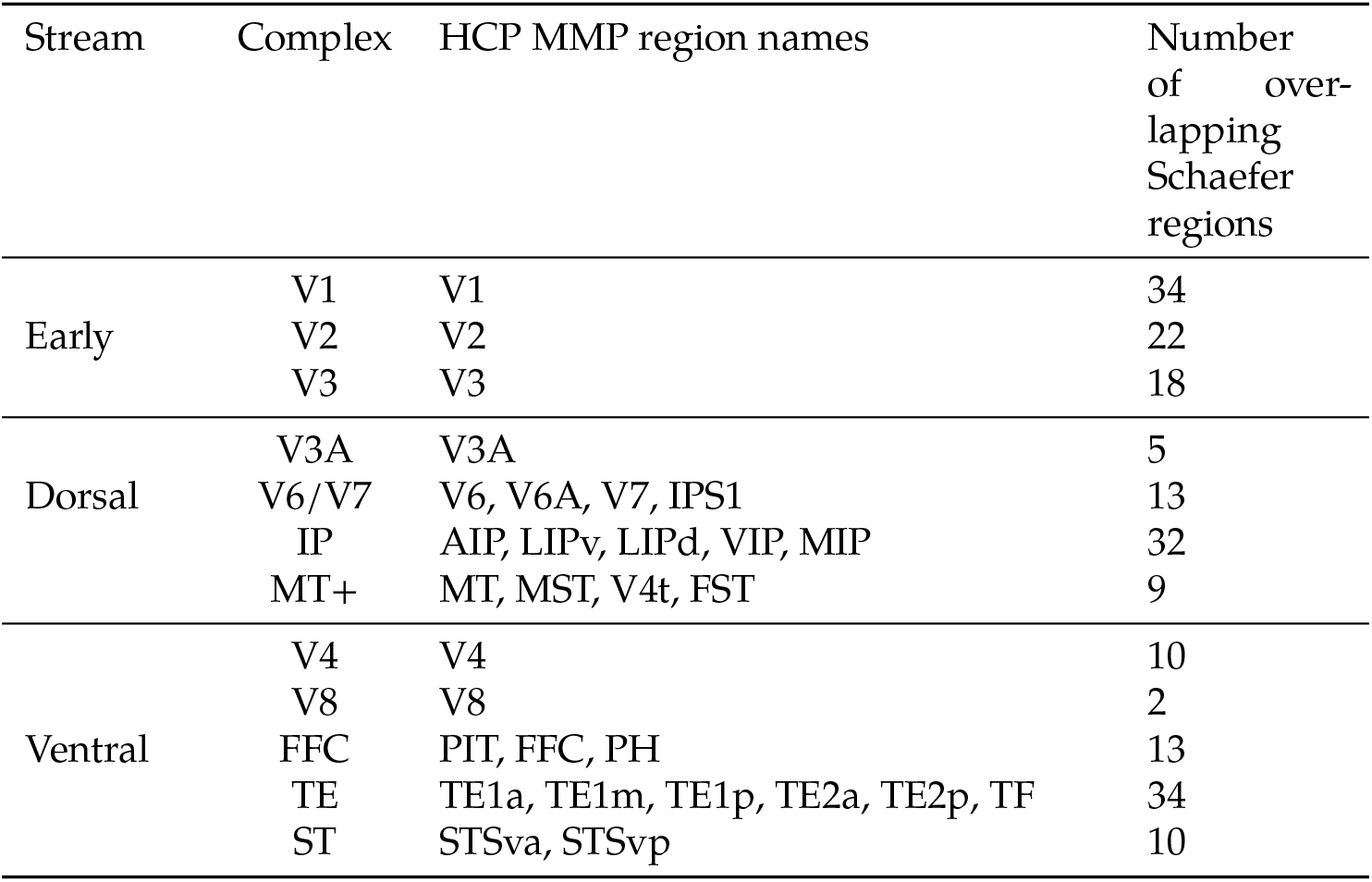
Parcellation of HCP MMP regions into complexes in the visual streams. Each visual stream consistes of multiple complexes, and each complex consists of one or more labels in the HCP MMP parcellation. Overlapping the HCP MMP and Schaefer1000 parcellations in fsaverage space assigns anatomical descriptions to Schaefer regions based on the most common within-region vertex label. IP, intraparietal complex; FFC, fusiform face complex; TE, inferior temporal complex; ST, superior temporal complex.

